# Social feedback structures stay/switch decision-making in *Drosophila*

**DOI:** 10.64898/2026.10.01.756036

**Authors:** Max J. Aragon, Mala Murthy

## Abstract

Within natural environments, different sensory stimuli can drive mutually-exclusive behaviors such as fighting, feeding, or courting. When experiencing competing cues, how do animals decide to continue executing the current behavior or to switch to a new behavior? To address this question, we developed a closed-loop behavioral paradigm that presents actively courting male *Drosophila* with a fictive heat stimulus, forcing the male to choose between two competing behaviors: continue courting (stay) or escape the heat (switch). By modeling the sensorimotor transformations underlying this decision process, we discovered that male flies continually evaluate social cues—including the female’s speed and her distance—in order to make a stay/switch decision. We investigated the neural correlates of stay/switch decision-making via brain-wide calcium imaging in head-fixed male flies presented with both fictive female visual cues and heat stimuli. We built models that predicted stay/switch decisions from neural data, which revealed that activity in a small number of brain regions, including the anterior optic tubercle, predicted decision outcomes up to 4 seconds prior to stimulus onset, whereas after stimulus onset, distributed activity across the brain was correlated with the behavioral decision. Together, these results demonstrate that *Drosophila* continually evaluate social feedback to inform decisions about how to respond to environmental cues, and that brain activity changes prior to presentation of the environmental cue can inform upcoming decisions. Overall, this study provides new insight into the processes underlying naturalistic decision-making in *Drosophila*.

## Introduction

Animals in natural environments face a constantly changing sensory landscape, requiring continual evaluation of which cues to prioritize and which to ignore, when multiple stimuli compete for attention and can potentially drive mutually exclusive behaviors. For example, a foraging animal might simultaneously encounter food odors, the presence of potential mates, and predator threats—each demanding different actions. Decision-making in such complex environments has attracted considerable attention across both neuroscience and ethology [1–7]. In flies, copulation persistence in the presence of aversive sensory stimuli depends on copulation duration: copulation persistence is high during early mating but decays over time [8, 9]. Similarly, courting male flies are more likely to ignore a looming predator-like visual cue if they have already invested significant time in courtship [10]. These studies suggest that the decision to persist with the current behavior or switch to a different behavior may depend on relatively simple decision rules, such as how long the animal has invested in the current behavior. However, *Drosophila* courtship is a highly dynamic process; for example, male courtship song production is continuously sculpted by changing female feedback [11] and fluctuating internal states [12]. These findings raise the possibility that persistor-switch decisions depend not only on time invested, but also on ongoing changes in the interaction itself—how well the male is performing the current behavior, and whether his social partner is responding to him. In other words, male flies may utilize ongoing feedback cues from their social partners to continually evaluate whether to persist with courtship or switch to another behavior in response to aversive stimuli.

This framing of decision-making draws from foraging theory, opportunity cost models, and action selection mechanisms [13–18], which posit that animals should evaluate both the value of their current activity and the potential benefits of alternative behaviors. However, the neural mechanisms that support such ongoing evaluations, particularly at the level of brain-wide neural dynamics, remain largely unexplored. Examining brain-wide activity is critical in this context, given that decision-making likely engages distributed circuits [19].

Here we investigate the mechanisms that shape ongoing decisions during courtship in *Drosophila melanogaster*. We developed a closed-loop behavioral paradigm that presents courting male flies with an aversive optogenetic heat stimulus. The heat stimulus is triggered by a specific syllable of the male’s song production. Despite providing the aversive cue during the same courtship context each time, we found optogenetic stimulation drives a mix of “stay” and “switch” responses, which are characterized by a continuation or termination of courtship, respectively. By tracking male and female behavior [20], we uncovered the social cues that influence this decision-making. Particularly, changes in female forward velocity, male-female distance, and the relative angle between flies in the moments prior to optogenetic stimulation predict the male’s decision to stay or switch, indicating the male’s continual evaluation of female behavior not only drives his choice of courtship behavior [11] but also his future decisions about responses to environmental cues. Brain-wide 2-photon imaging during the simultaneous presentation of female stimuli and fictive heat revealed a number of brain regions with decision-predictive activity several seconds prior to stimulation. In addition, we uncovered even more neuropils with post-stimulus activity that were predictive of stay versus switch. Altogether, our results suggest that animals dynamically evaluate the state of their current behavior in order to decide whether or not to prioritize novel sensory cues, and that coordinated dynamics across the brain support this decision-making process.

## Results

### *Drosophila* males make stay/switch decisions during courtship

During courtship, males pursue females and sing to them [11, 21]. To generate an aversive stimulus during courtship, we chose to optogenetically activate hot cells (HCs), a small population of thermosensory neurons in the aristae that encode noxious heat [22] and drive escape behaviors [23]. We first identified the minimal optogenetic stimulus that reliably elicits escape behaviors in solitary male flies (Fig. SI1A; see Methods), including increases in bilateral wing spreading and increases in speed (Fig. SI1B; Supp. Movie 1). These responses likely comprise a take-off escape response similar to that observed in the context of a visual loom threat [24, 25].

Using this minimal stimulus, we asked how the male fly responds to HC stimulation during the courtship context. We triggered HC stimulation on the production of male sine song, using a novel neural network that detects sine song in real time (Fig. 1A; see Methods). We targeted this song syllable because males consistently produce sine song near the female [26]. To prevent male flies from learning the relationship between sine song and the stimulus, our stimulation protocol used a two-stage design. During the first “unlocking stage,” sine song detection triggers a five second timer during which song production does not result in optogenetic stimulation. After the timer elapses, subsequent sine song detection leads to a 200 ms red LED pulse that activates csChrimson [27] in HCs in the male (female flies were wildtype and did not respond to the stimulus). Empirically, the median inter-stimulus interval (ISI) across our dataset was 33.5 seconds (Fig. SI1C).

**Fig. 1.**
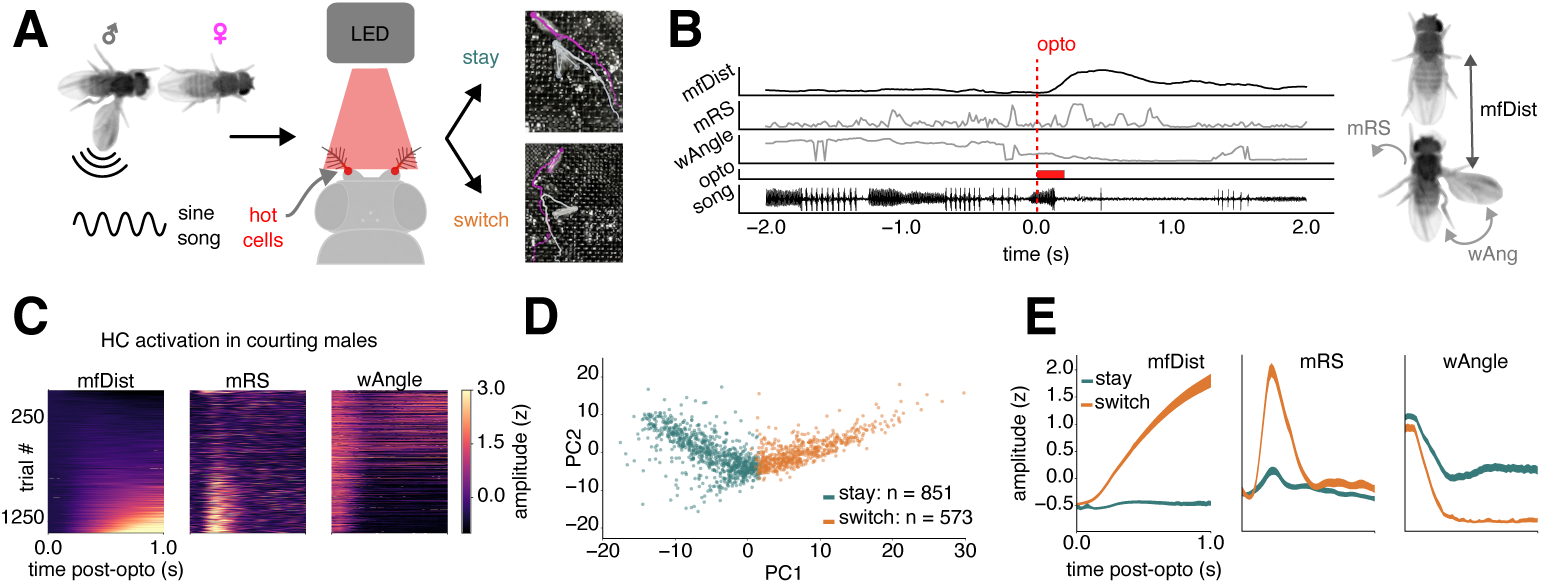
*Drosophila* males make stay-switch decisions during courtship. (A) During courtship, sine song production (left) triggers optogenetic hot cell activation (middle), which results in either “stay” and “switch” decisions (right) in male flies expressing the light-sensitive opsin. (B) Example of a decision-making trial during courtship. Left: behavioral features (male-female distance (mfDist), male rotational speed (mRS), male unilateral wing angle (wAngle), and song) aligned to onset of the optogenetic stimulus during one example trial. Right: illustration of mfDist, mRS, and wAngle. (C) Stimulus aligned mfDist (left), mRS (middle), and wAngle (right) across all trials (n=46 experiments, n=1424 trials) sorted by average mfDist. (D) Two-dimensional principal components embeddings of mfDist, mRS, and wAngle traces concatenated for each trial. Stay trials (teal) and switch trials (orange) are clustered using a two-cluster K-means algorithm. (E) Cluster-averaged stay (teal) and switch (orange) traces aligned to the optogenetic stimulus for mfDist (left), mRS (middle), and wAngle (right). Bands indicate 95% confidence intervals.

To verify that males did not learn the relationship between song production and opto stimulation, we tested the dependence of ISI on the number of stimulation events, with the rationale that learning would increase this interval over time in order to minimize aversive heat stimulation. We fit a linear-mixed effects model (LMM) that predicts the interval between stimulus-producing song bouts from the number of stimulation events, and found no significant relationship effect (p = 0.077; see Methods). This suggests that flies did not modify the frequency of song bouts in response to repeated HC stimulation. Altogether, this experimental design allowed us to precisely examine, on a trial-by-trial basis, whether males respond to or ignore the noxious cue during courtship, with each trial defined as the time window surrounding an optogenetic stimulus (Fig. 1B).

When we activated HCs in courting males (n=46 pairs, n=1424 trials, trials per experiment = 31 *±* 26), escape behaviors appeared to be more subtle than in the solitary context (Supp. Movie 2). During courtship, we found changes in three behavioral features relative to HC stimulation: increases in male-female distance (mfDist) and male rotational speed (mRS), and decreases in the angle of the song-producing wing (wAngle) (Fig. 1B-C). These behaviors are consistent with males terminating courtship. However, we observed variability in these behavioral responses, suggesting that males only sometimes terminated courtship.

To investigate this variability, we clustered the stimulus-aligned mfDist, mRS, and unilateral wAngle responses using PCA and K-means clustering (Fig. 1D). We found that a clustering assignment with two trial types was the most parsimonious description of our data (Fig. SI1D). Trial embeddings were separable along the first principal component, the loadings of which are associated with a sustained increase in mfDist, a pronounced yet transient peak in mRS, and a decrease in wAngle (Fig. SI1E). Trial-averaged behavior for cluster 1 showed little stimulus-induced change in these variables, while cluster 2 showed pronounced effects (Fig. 1E). We labeled trials within the first cluster as “stay” trials because the male continued producing courtship behaviors—including extending his song-producing wing and staying close to the female—and labeled trials within the second cluster “switch” trials because the male switched from close courtship to threat evasion (in particular, U-turns are characteristic of heat avoidance [23]).

Flies tended to show a mix of stay and switch trials throughout an experiment (fraction of switch trials = 0.46 *±* 0.26), rather than only expressing one decision outcome (Fig. SI1F). In addition, the probability of stay or switch did not consistently change as a function of time (Fig. SI1G; black line indicates empirical switch probability), meaning that males flip back and forth between stay and switch decisions rather than changing their behavior as they invest more time in courtship. This contrasts with recent work on visual threat-driven escape responses during courtship [10] but is more similar to alternations between different internal states across time during courtship [12]. We also tested whether the inter-stimulus interval (ISI) predicts stay/switch outcome, as we reasoned that more frequent HC activations could bias the animal’s decision strategy. We found that longer, not shorter, ISIs predicted switch decisions (logistic regression, *p* = 0.007; see Methods), which is opposite to the prediction of a stimulus carryover effect.

### Drosophila males continuously monitor female feedback to inform stay versus switch decisions

We next investigated whether cues provided by the female during courtship impact stay (continue courtship) versus switch (suspend courtship) decisions. We built on previous work investigating the relationship between social feedback and song patterning [11, 12] to model the sensorimotor transformations underlying stay/switch decisions. We quantified 16 movement and interaction features (Fig. 2A), including female movements (female forward velocity (fFV), female forward acceleration (fFA), female rotational speed (fRS), female lateral speed (fLS), and female lateral acceleration (fLA)), male movements (male forward velocity (mFV), male forward acceleration (mFA), male rotational speed (mRS), male lateral speed (mLS), and male lateral acceleration (mLA)), male-female distance (mfDist), male-female angles (mfAngle and fmAngle), times when the male taps the female abdomen (taps) [26], times when his head is positioned underneath her wings (under wings; likely corresponding to proboscis licking) [28], and ongoing experiment time. We then fit generalized linear models (GLMs [11]) using five seconds of feature history to determine if and how these features predict decision outcomes (Fig. 2B; see Methods). We found that female forward velocity (fFV) was the most predictive of stay/switch outcome (Fig. 2C). We then examined which additional features, when combined with fFV, increased stay/switch predictive power by at least 2% on average. We found that four additional features—mfDist, fmAngle, mfAngle, and mRS—satisfied this criterion (Fig. 2D).

**Fig. 2.**
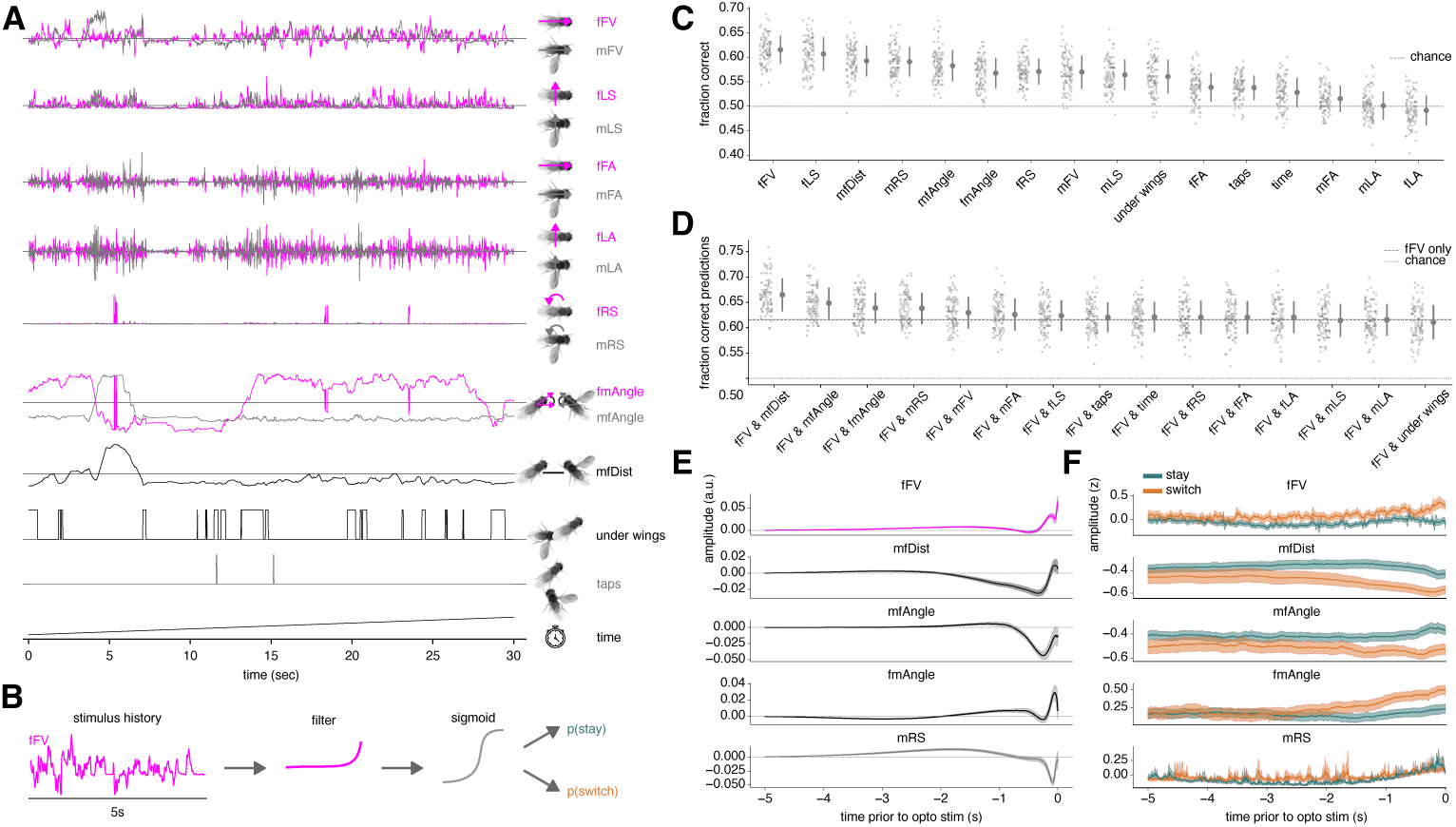
Changes in social feedback predict stay/switch decisions. (A) 16 social cues used in our model, including: female forward velocity (fFV) and male forward velocity (mFV); female lateral speed (fLS) and male lateral speed (mLS); female acceleration (fFA) and male forward acceleration (mFA); female lateral acceleration (fLA) and male lateral acceleration (mLA); female rotational speed (fRS) and male rotational speed (mRS); female-male angle (fmAngle) and male-female angle (mfAngle); male-female distance (mfDist); under wings; taps; and time. Female-specific features are colored magenta, male-specific features are colored gray, and relative features are colored black. (B) Generalized linear modeling (GLM) pipeline. Five seconds of stimulus history are convolved with a linear filter, the output of which is passed through a nonlinearity to generate a probability of stay versus switch. (C) Fraction of correct predictions on held-out data for GLMs fit with a single feature. Features are ordered left to right from most predictive to least predictive. For each feature, the full distribution of held-out scores is shown (each dot indicates performance on a cross-validation fold) in addition to the mean and standard deviation of scores across folds (see Methods). (D) Same as (C), but for GLMs fit with both fFV (the feature with greatest stay/switch predictive power alone) and one additional feature as input. (E) GLM filters (mean and standard deviation across cross validation folds) for fFV and features that contributed at least 2% average increase in fraction of correct predictions across cross validation folds. (F) Five seconds of z-scored behavioral features preceding optogenetic stimulation (mean and 95% confidence intervals), grouped by stay (teal) and switch (orange) trial outcomes.

Next, we examined the linear filters associated with our two-feature GLM models, as these filters reflect how the male fly integrates cues over time to guide decision-making. The fFV filter ramps up sharply in the *∼*200ms before stimulus onset (Fig. 2E, top row), indicating that high female forward velocity in this window predicts a switch decision. The mfDist filter is biphasic (Fig. 2E, second row): a negative lobe emerging *∼*2s before stimulation—consistent with the short inter-fly distance typical of sine song—followed by a sharp positive lobe in the final *∼*200ms, reflecting the female moving away from the male just before he switches. mfAngle (the female’s position relative to the male’s heading) develops a negative lobe beginning *∼*1s before stimulation (Fig. 2E, third row), while fmAngle (the male’s position relative to the female’s heading) develops a positive lobe over roughly the same window (Fig. 2E, fourth row). Together with the rise in mfDist, this pattern is consistent with the female moving away along a path that remains within the male’s heading axis (small mfAngle) while he falls off her heading axis (large fmAngle)—as would occur if he does not reorient quickly enough to keep pace with her departure. Finally, the mRS filter shows a brief negative lobe just before stimulation (Fig. 2E, bottom row). Altogether, these filters suggest that switch trials are preceded by the female speeding up and moving away from the male.

To confirm our interpretation of these GLM filters, we examined raw feature traces prior to stimulus onset sorted by stay and switch trials. Four features—fFV, mfDist, mfAngle, and fmAngle—showed a clear separation between conditions consistent with their GLM weights (Fig. 2F). mRS, by contrast, showed no clear trial-averaged separation despite carrying predictive weight in the model. This dissociation suggests that mRS itself may not carry direct predictive information about stay/switch outcome; rather, its inclusion in the GLM may reduce shared noise variance in other features.

Altogether, our GLM analysis indicates that the male continually evaluates female speed, distance, and relative position, and uses this information to decide whether to continue with courtship or abandon courtship in response to a future threat cue. Importantly, we find that the male evaluates these cues *independent* of whether the heat stimulus is provided, since the optogenetic stimulus is not delivered at predictable time intervals (Fig. SI1C).

### Flies make stay/switch decisions during brain-wide imaging

How does the brain support stay/switch decision-making during courtship? To address this question, we designed an experiment in which we presented a head-fixed and walking male fly with a moving virtual female object projected onto a spherical display during 2-photon imaging (Fig. 3A). To induce heat-evoked turning behavior in this setup, we integrated a red LED into the objective for optogenetic stimulation of HCs. The stimulation strength closely matched our free behavior experiments (see Methods). We presented two cohorts of flies with a virtual female target on the right visual field. For one cohort (female +), we also adhered a live female fly to the recording platform on the male’s left side so the male could tap her to induce a courtship state [29] (Fig. 3B, left). For the other cohort (female -), we presented only the virtual female without a tethered, live female (Fig. 3B, right). In both cases, the male’s left eye was blinded (to eliminate conflicting visual cues from the live female in the female + case; see Methods). We expressed the calcium sensor GCaMP6s [30] in all neurons using the nsyb genetic driver [31, 32], which allowed us to sample neural dynamics broadly throughout the brain [33]. We simultaneously recorded the male’s behavior, and related both to the motion of the virtual female object (Fig. 3C). To facilitate cross-session analysis of neural activity, we segmented neural data at the level of supervoxels—spatially contiguous voxels with correlated activity (Fig. 3D, top)—and registered these functional units into the Functional Drosophila Atlas (FDA) anatomical space using the BIFROST registration pipeline (Fig. 3D, bottom) [34], which permitted subsequent analysis of neural activity at the level of individual neuropils.

**Fig. 3.**
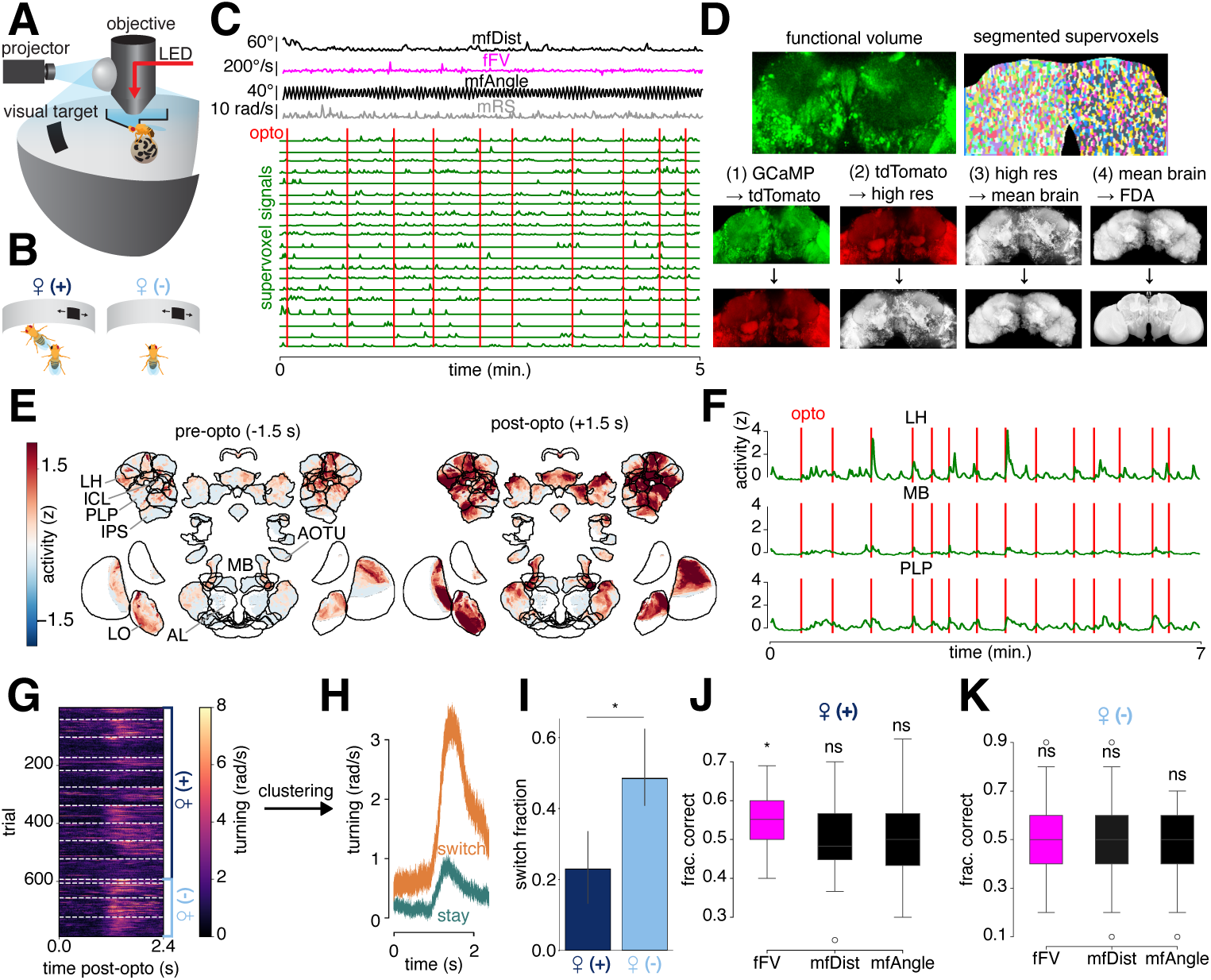
Flies make stay/switch decisions during brain-wide 2-photon imaging. (A) Experimental setup for uncovering neural signatures of stay/switch decisions. (B) Top down view of male on 2-photon rig. The female + cohort (left) experienced a moving visual target and cues from a tethered, live female. The female - cohort (right) experienced the visual target but not the live female. In both cases, the male’s left eye was painted to render him blind in the left eye (so the visual target could only be seen on the right). (C) Example of visual stimulus dynamics (mfDist, fFV, mfAngle), male speed (mRS), and neural activity (green traces) recorded during each experiment (see Methods). (D) Pipeline for supervoxel segmentation (top row) and functional volume alignment to the Functional Drosophila Atlas (bottom row). (E) Example of brain-wide activity from one female + animal before optogenetic stimulation of HCs (left) and after stimulation (right). (F) Neural activity in neuropils implicated in thermosensory processing, including the lateral horn (LH), mushroom body (MB), and posterior-lateral protocerebrum (PLP). Optogenetic stimulation of hot cells is indicated by vertical red lines. (G) Male rotational speed triggered on optogenetic stimulation. White dashed lines delineate different experiments. Female + experiments are indicated by the dark blue bracket; female - experiments are indicated by the light blue bracket. (H) A two-cluster K-means clustering algorithm partitioned opto-aligned behavior into stay-like turning behavior (teal trace) and switch-like turning behavior (orange trace). Shaded regions indicate 95% confidence intervals. (I) Comparison of switch trial fractions between experiments with a female fly (dark blue) and without a female fly (light blue). (J) Fraction of correct predictions from a GLM fit on fFV, mfDist, and mfAngle cues from the virtua_1_l_0_female in the female + context. Magenta indicates female-specific cues, while black indicates relative cues. (K) Same as (J), but for the female - context.

In response to optogenetic HC activation (see Methods for optogenetic activation protocol in head-fixed flies), we observed widespread responses in the brain (Fig. 3E), including in neuropils implicated in temperature coding such as the lateral horn (LH), mushroom body (MB), and posterior-lateral protocerebrum (PLP) (Fig. 3F) [22]. This stimulus-induced activity was not a response to the light flash alone or due to changes in walking behavior (Fig. SI2A-B). In the seconds following optogenetic stimulation, we observed variable, stimulus-induced behavioral responses in ATR+ males (males with active opsin), with some trials showing little opto stim-induced turning and some trials showing large turning responses (Fig. 3G). This was similar to free behavior, in which we observed a mix of stay/switch trials (Fig. 1). Our ATR- controls (inactive opsin) showed no obvious turning in response to the optogenetic stimulus (Fig. SI2C). Does the behavioral variability we observe in the ATR+ group reflect stay/switch decision-making? We reasoned that true stay/switch decisions must satisfy three criteria. First, rotational speed should cluster into two distinct trial types, with one cluster showing little or no turning (stay trial) and the other cluster showing robust turning (switch trial). Second, the probability of observing a stay or switch trial should depend on the presence of a live female fly—specifically, courtship state should bias the animal toward making stay decisions. Finally, stay/switch decisions should be predictable from fictive social cues (movement of the female). We found that the male’s behavior did cluster into two response types: average rotational speed within each cluster showed a stay-like (teal) and switch-like (orange) response (Fig. 3H). We also found that compared to the female + cohort, the female - cohort made proportionally more switch-like decisions (Fig. 3I). Finally, we constructed GLMs that predict stay/switch outcomes from the visual cue, including female forward velocity (fFV) and male-female distance (mfDist; corresponding to the size of the visual cue). In the female + context, we found that only fFV predicted stay/switch outcomes above chance (Fig. 3K). In contrast, no features in the female - context predicted stay/switch outcomes above chance (Fig. 3L). Altogether, our head-fixed flies satisfied all three criteria, suggesting that they produce stay/switch decisions.

### Neural dynamics predict stay/switch

We next investigated whether neural activity predicts stay-switch decision outcomes; imaging pan-neuronally enabled us to search for such activity anywhere in the brain. Supervoxels (Fig. 3D) could be averaged to examine neuropil-level responses (Fig. 3F; see Methods). We fit GLMs (similar to Fig. 2) that convolve neuropil-averaged activity with a linear filter to predict stay/switch outcome (degree of turning on the ball, (Fig. 3H) either before or after the presentation of the optogenetic stimulus (Fig. 4A; Methods). We found that ten neuropils contained decision-predictive activity *before* stimulus onset (Fig. 4B), the most predictive of which was the anterior optic tubercle (AOTU), a brain area previously shown to contain neurons that are both modulated by male courtship state and that represent female motion cues [21, 35]. When we examined neural activity after stimulus onset that predicts stay/switch decisions, we found that 28 different neuropils predicted decision outcome, including the inferior clamp (ICL), inferior posterior slope (IPS), superior posterior slope (SPS), and posterior lateral protocerebrum (PLP) (Fig. 4C). Neuropil activity before stimulus onset that was predictive of stay/switch decisions were largely concentrated in the SEZ and along midline structures (Fig. 4D, top), whereas neuropil activity after stimulus onset that was predictive of stay/switch decisions was broadly distributed across the brain, including most protocerebral regions (Fig. 4D, bottom).

**Fig. 4.**
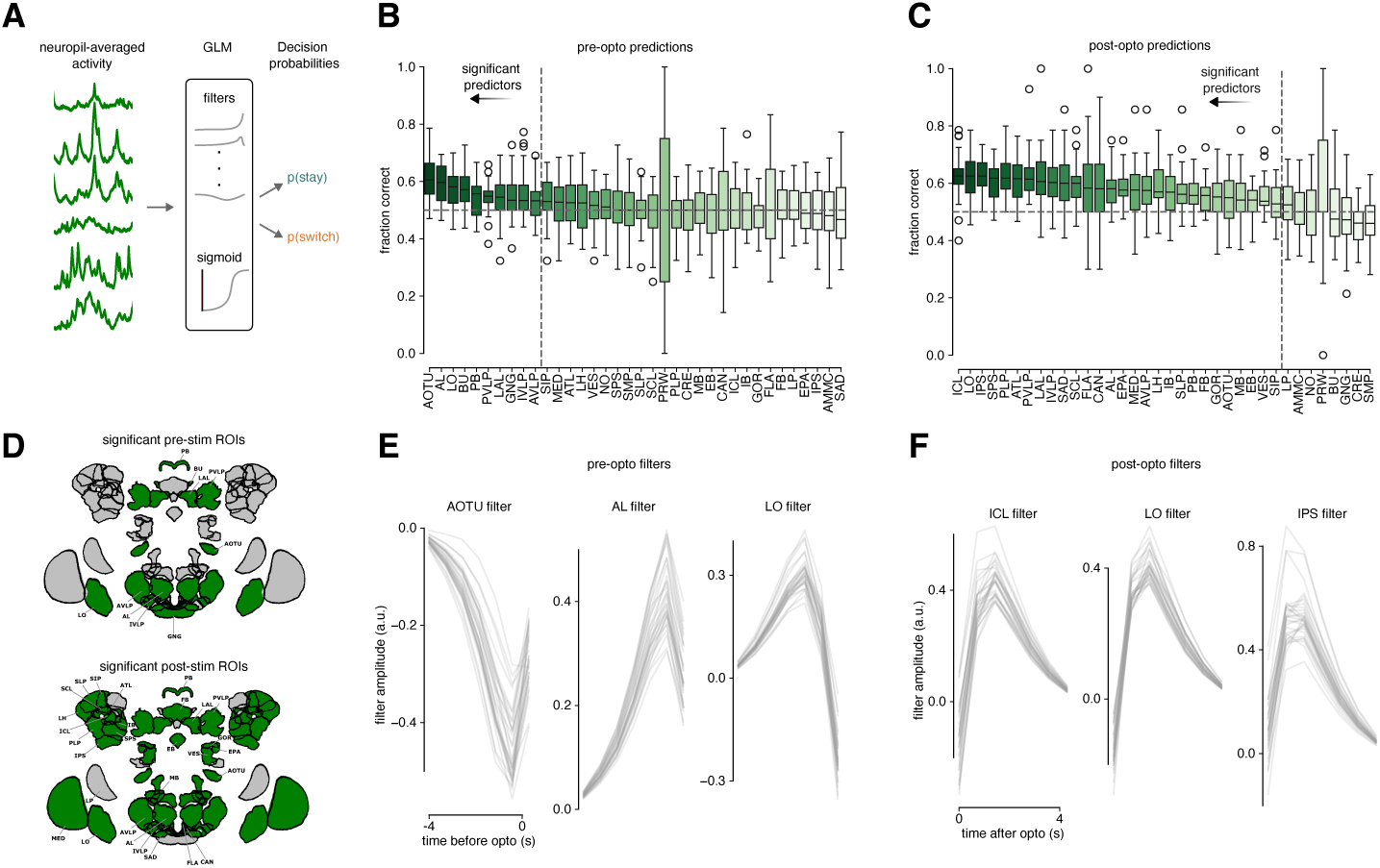
Neural dynamics predict stay/switch decisions. (A) GLM modeling strategy. Neuropil-averaged activity preceding or following optogenetic stimulation was passed through GLM filtering and nonlinearity stages to generate a stay/switch probability. (B) Fraction of correct predictions on heldout data for GLMs trained to predict stay/switch outcome from neuropil-averaged activity preceding the optogenetic stimulus. Neuropils are sorted by median performance. Vertical dashed line indicates significance threshold after correcting for multiple comparisons. Horizontal dashed line indicates chance performance. (C) Same as (B), but for GLMs trained on neural activity after the optogenetic stimulation. (D) Neuropils with decision-predictive activity (green) when using neural activity preceding the optogenetic stimulus (top) or following the optogenetic stimulus (bottom). (E) GLM filters for the top three neuropils that best predict stay/switch outcome from dynamics preceding the optogenetic stimulus. (F) Same as (E), but for neuropils that best predict stay/switch outcome from dynamics following the optogenetic stimulus.

Next, we examined the models’ GLM filters to understand how neural dynamics translate to decision outcomes. Starting with pre-stimulus decisions, the GLM filter for the AOTU showed a negative peak prior to stimulus onset, indicating that lower activity in this brain region predicts switch decisions (Fig. 4E, left). This is consistent with increases in activity in AOTU neurons underlying courtship pursuit [21]; in other words, lower activity in the AOTU would relate to less engagement with the female, and increase the probability of a ‘switch’ decision. Interestingly, the filter shapes for the next most highly predictive regions, the antennal lobe (AL) and lobula (LO), showed different activation patterns. For the AL, the posterior lobe of which is involved in thermosensory processing [22], the filter had a single positive lobe (Fig. 4E, middle), indicating higher activity in this region prior to opto stimulus predicts switch decisions; in other words, fluctuations in neural activity outside of the courtship pathway may bias males towards switch decisions. The filter for the LO, on the other hand, showed a more complex biphasic filter (Fig. 4E, right). For post-stimulus activity, the three most predictive GLM filters peaked shortly after stimulus onset and all contained positive integrals, indicating that higher stimulus-evoked activity in these brain regions predicts switch trials (Fig. 4F).

Altogether, these results suggest that several seconds prior to optogenetic stimulation, activity in a specific set of brain regions, including visually-selective (AOTU and LO) and thermosensory (AL) neuropils, can predict the male fly’s decision to continue or terminate courtship. In contrast, the majority of neuropils in the brain contain decision-predictive activity after the presentation of the stimulus, consistent with switch decisions engaging distributed sensorimotor circuits across the brain.

## Discussion

We developed a closed-loop behavioral paradigm to investigate how male flies respond to an aversive environmental cue during active courtship, with the goal of understanding how animals evaluate new stimuli relative to ongoing social interactions. We found that the aversive stimulus elicits a variable response: either a “stay” decision where the male continues to court the female or a “switch” decision where the male abandons courtship by performing an evasive U-turn. Through behavioral modeling (GLMs), we uncovered that male flies continuously evaluate their interactions with the female—by keeping track of female forward velocity, male-female distance, and the angle between the male and female—in order to decide to either stay or switch. Furthermore, our GLMs revealed internally consistent sensorimotor transformations: males decide to switch when the female speeds up and moves away prior to hot cell (HC) stimulation. We also showed that during 2-photon imaging, head-fixed males show variation in stay/switch decisions following HC activation, decisions that depend on the behavior of a virtual female. By modeling the relationship between stay/switch outcome and neural dynamics on a neuropil-by-neuropil level, we discovered that a relatively small set of brain regions, including the AOTU, AL, and LO, contain decision-predictive information prior to activation of HCs, while activity after HC activation, in a broad array of neuropils, is predictive of stay/switch outcomes. We thus map brain-wide dynamics underlying stay/switch decision-making in *Drosophila*.

Several prior studies have explored behavioral strategies and neural circuits underlying how animals decide between multiple competing demands, such as the choice to satisfy hunger or thirst [7] or whether to avoid a predator-like cue or court a female fly [10]. Our study builds on this literature by asking whether animals can solve this problem by evaluating information about their current task. The discovery that social feedback dynamically shapes the male’s decision to continue courting in the presence of an aversive environmental cue provides evidence in favor of this idea, and provides a solution to how flies constrain the vast space of possible behaviors.

Why might evaluating social cues, such as the female’s forward velocity, be an adaptive strategy for solving the stay/switch problem? Previous work shows that sexually receptive female flies tend to slow in response to courtship song [36, 37]. In our experiments, optogenetic stimulation of HCs was yoked to male sine song production; during the time preceding stimulation, the male may treat the female’s choice to slow down to his song as a receptivity cue, with increases in speed indicative of sexual rejection. Two mechanisms could explain this relationship: rejection signals could actively prime the male to attend to threats from predators, heat sources, or other flies, or alternatively, female receptivity cues could suppress attention to non-courtship stimuli, with their withdrawal releasing that suppression. Our neural data are consistent with both mechanisms: decreased AOTU activity before switch decisions suggests a release of courtship-driven engagement, while increased AL activity may reflect heightened sensitivity of thermosensory circuits. Distinguishing between these possibilities—active priming versus release from inhibition—remains an important direction for future work. Regardless of mechanism, the influence of social feedback on threat responsiveness is a natural extension of Tinbergen’s idea of “sign stimuli”—particularly stimuli that drive behavioral patterns [38]—applied to the transitions between orthogonal behaviors such as courtship and threat evasion.

Our experiments investigating brain-wide neural dynamics during a virtual courtship interaction provide insight into how the brain mediates this sophisticated decision-making process. Such a brain-wide screen was critical as we did not know, a priori, which neuropils or cell types would contain decision-predictive activity. We showed that dynamics within the AOTU prior to heat stimulus onset can predict stay/switch outcomes (Fig. 4B), and that activity in this region tends to decrease during switch trials (Fig. 4E, left). The AOTU is densely innervated by LC10a visual projection neurons and downstream targets, which all encode the motion of small objects [35, 39]. Importantly, LC10a activity is state-dependent: during periods of high courtship motivation, these neurons respond robustly to female-like stimuli, and become quiescent during periods of low motivation [21]. Our finding that AOTU activity decreases prior to switch decisions is consistent with this finding: the male may enter a disengaged state prior to a switch decision, which is reflected in low AOTU neuron activity.

Interestingly, two other neuropils with predictive power before stimulus onset—the AL and LO—showed different activation patterns compared to the AOTU. The AL is a component of the thermosensory pathway, and increased activity in this region preceding switch decisions (Fig. 4E, middle) could reflect a disengagement state-related increase in basal activity. This increased activity is unlikely to be residual from the previous optogenetic stimulation, as the minimum inter-stimulus interval in our experiments (15 seconds) far exceeds the timescale of stimulus-evoked responses in these circuits. Instead, this state-dependent change could, in principle, sensitize the animal to subsequent thermosensory cues and hence promote a robust escape response. The LO, in contrast, showed biphasic activity (Fig. 4E, right), indicating that the temporal dynamics of neural activity, rather than its overall level, within this region predict stay/switch. This pattern may represent the LO detecting changes in the size of the female on the male retina, specifically when this size decreases as she moves away from the male. Together, the AL and LO may therefore coordinate, through a currently unknown circuit mechanism, to promote threat-evoked courtship termination in the context of social cues that signal sexual rejection.

Our ability to predict stay/switch decisions from neural activity in a restricted number of neuropils has parallels in recent work examining volitional locomotor control. Brezovec et al. [40] performed brain-wide imaging in walking flies and found that activity in the inferior posterior slope (IPS) could predict future turning direction, even in the absence of any external sensory cue—suggesting that upcoming, internally generated actions are encoded in brain-wide activity patterns well before their execution. Our results extend this principle to a distinct behavioral context: we show that pre-stimulus activity in a small set of neuropils shapes how the animal will respond to a subsequent sensory cue (optogenetic activation of HCs). Together, these findings suggest that the fly brain maintains a representation of upcoming behavioral choices, whether those choices are volitionally generated or in response to external stimuli.

How does the male’s motivation to court allow him to disregard environmental threats during stay decisions? Our finding that many neuropils’ post-stimulus dynamics predict stay/switch outcomes suggests that this “tunnel vision”-like state involves coordinated activity across the brain. Presumably, thermosensory information is represented differently during a stay and switch trial. How this information is transformed—whether through gain modulation or a more nuanced mechanism at the level of population dynamics—remains open, but addressing it will require recordings from a variety of cell types across the brain.

In conclusion, continual evaluation of social feedback provides a solution to the problem of choosing one among many possible behaviors. We find that flies make principled decisions about when to persist and when to switch on a moment-to-moment basis. Our findings provide a framework for uncovering how the brain supports context-appropriate actions in a world full of choices.

## Author Contributions

M.J.A and M.M. designed the study. M.J.A. collected, processed, and analyzed the data. M.J.A. and M.M. wrote the paper. M.M. obtained funding.

## Acknowledgments and Funding

We thank Stephan Thiberge, Albert Lin, Bella Brezovec, and Megan Wang for assistance in establishing the two-photon imaging system and pipeline for analysis of calcium imaging data, Anthony Crown for design of Fig. 3A-B, and Ayelet Sarel and members of the Murthy lab for comments on the manuscript. This project was supported by the following grants to MM: NIH NINDS Research Program Award (R35) and NIH BRAIN Initiative Award (R01 NS104899).

## Methods

### Fly husbandry and stocks

Virgin male flies were single-housed for 4-7 days and stored in a 25C incubator prior to imaging. 0-7 day old virgin females were group housed in low density (*≤* 4 animals per vial). All animals were fed a standard diet except those fed ATR food. Fly stocks included the following: **(1)** *NM* 91; **(2)** *w*^1118^, *20xUAS-Chrimson*; *HC-Gal4* /+; +; **(3)** *w*^+^/Y; *HC-Gal4* /*lexA-nsyb*; *20xUAS-Chrimson*/*LexAop-GCaMP6s*, *tdTomato*

### Behavioral chambers

We used the same behavioral chambers described in Roemschied et al., 2023 for our stay/switch assay [26]. Briefly, behavioral experiments were conducted in two custom circular chambers housed within black acrylic enclosures, illuminated by white LED pads (Metaphase Technologies). Video was recorded at 60 fps using FLIR Blackfly S cameras (BFS-U3-13Y3M-C, 25 mm TechSpec VIS-NIR lens) with infrared illumination ( 22 *µW/mm*^2^) and an IR bandpass filter (850 nm CWL, 10 nm FWHM, Thorlabs) to block optogenetic red light. Recordings were made via the Motif system (loopbio GmbH). Sound was captured at 10 kHz from 16 particle velocity microphones (Knowles NR-23158-000) embedded in the chamber floor and coated with IR-absorbing dye (Epolin Spectre 160) to minimize video artifacts. Chamber temperature was monitored using an analog thermosensor (Adafruit TMP36).

### Optogenetics

Flies were kept for 3-5 days on food supplemented with all-trans retinal (ATR) at 1 ml ATR solution (100 mM in 95% ethanol) per 100 mL of food. ATR-fed flies were reared in the dark. We identified the minimal optogenetic stimulus for eliciting robust behavioral responses in solitary males by randomly presenting animals with a stimulus sweep in the range 1-205 *µW* mm*^−^*^2^ with a pulse duration in the range 0.05-1.0 seconds. For all subsequent free-behavior experiments with solitary and courting males, we used a stimulus with 0.544mW*/*cm^2^ intensity and 200 ms duration. This step was critical because over-activation of the thermosensory pathway could override the modulatory effects of social feedback on the male’s behavior during courtship. We tested various combinations of stimulation duration and LED driver voltage, and found that a 0.05 V stimulus lasting 200 ms reliably evoked behavioral responses. When we presented solitary flies (n=24) with this stimulus, we observed consistent bilateral wing spreading and sharp increases in forward speed locked to optogenetic stimulus onset (Fig. SI1B).

### Closed-loop stay/switch assay

#### Online song segmentation

To enable real-time sine song detection, the data acquisition (DAQ) system delivered 8 ms of 16-channel audio per readout. Each 38.4 ms (384 sample) window was preprocessed as follows: we selected the channel containing the highest-power sample and used only that channel for further processing. The waveform was normalized by dividing it by its vector norm, then Fourier-transformed using 512 sampling points to extract the magnitudes of the signal’s frequency components. The frequency-domain data was then passed to a convolutional neural network (CNN).

The CNN model, which was implemented using Keras with Theano backend, consisted of two convolutional layers. The first layer applied eight 9-sample-wide filters with a stride of 1 and ReLU activation. The second layer used eight 9-sample filters with a stride of 2, followed by ReLU activation and max pooling with a pool width of 2. The output was flattened and passed through a dense ReLU-activated layer with 32 neurons, followed by a sigmoid-activated output neuron representing the probability that the waveform contained sine song. Each input produced a value between 0 and 1; values above 0.995 were classified as sine.

Training data were derived from 19 recordings of wild-type NM91 male-female courtship sessions. Windows were labeled as “sine” or “no-sine”, using outputs from an offline song segmenter as ground truth. Recordings were split into training (11), validation (4), and test (4) sets. To address class imbalance, weighted losses were used during training so that a naive classifier would assign equal total loss to both classes. During training, a 25% dropout was applied to reduce overfitting.

#### Stimulation protocol

Sine-song detection triggered a five second timer during which sine song production did not result in optogenetic stimulation. The end of this five-second period unlocked the stimulation: subsequent sine song resulted in a 200 ms LED pulse.

#### Stay/switch decision assignment

For each trial, one second of post-stimulus mfDist, mRS, and wAngle were concatenated into a single vector, yielding a matrix of size 1424 x 180 (#trials x time). Next, we used PCA to reduce the dimensionality of these data to 19 dimensions, which accounted for 90% of the variance. We determined the optimal number of clusters (2) by calculating the silhouette score across a range of clusters (minimum: 2, maximum: 9). In our two-cluster partitioning of the behavioral data, we assign the cluster with higher average rotational speed as the “switch” type and the cluster with lower average rotational speed as the “stay” type.

#### Linear mixed-effects modeling to test heat conditioning

We fit a linear mixed-effects model (LMM) that predicts the inter-stimulus interval (ISI) between HC stimulation events from the number of stimulation events the animal received. If the animal learned the association, we reasoned that the ISI should increase as a function of stimulation events, reflecting increasing hesitance to sing to the female in order to minimize aversive heat stimulation. Importantly, the model leverages fly-specific relationships between ISI and stimulation number (“random effects”) in order to estimate a global effect shared across flies (“fixed effect”). The model was fit using the statsmodels package. The fixed effects formula included a trial number regressor, and the random effects formula included a random intercept term and a trial number regressor.

#### ISI–switch probability analysis

To test whether the inter-stimulus interval (ISI) predicts stay/switch outcome, we computed the ISI preceding each trial (excluding the first trial of each session, which has no preceding stimulus). ISIs above the 99th percentile were excluded to remove extreme outliers. We then fit a logistic regression model predicting stay/switch outcome from z-scored ISI using scikit-learn (no regularization penalty). Statistical significance was assessed via a two-sided permutation test (10,000 permutations), in which the stay/switch labels were randomly shuffled and the logistic regression coefficient was recomputed for each permutation. The *p*-value was defined as the fraction of permuted coefficients with absolute value greater than or equal to the observed coefficient.

### Generalized linear modeling

#### Predicting stay/switch during natural courtship

For free behavior experiments, we extracted five seconds of feedback cue history for each feature of interest, which yielded our design matrix. We then projected this matrix onto a filter bank consisting of 12 raised cosine bumps [41], which imposed a smoothness constraint on our filters while also reducing the number of learnable parameters to 13 (12 filter coefficients and an intercept term). Next, we randomly partitioned our data into training and validation sets (80% training, 20% validation) 100 times to assess how the models performed on held-out data. To ensure the stay/switch classes were balanced, we applied random oversampling for the training data. We also randomly under-sampled our validation data to achieve balanced classes, which allowed us to compare model performance to chance predictions. For model fitting, we applied L2 regularization with an inverse regularization strength of *C* = 3 *·* 10*^−^*^3^. Once we fit the GLMs, we reconstructed time-domain filters by back-projecting the GLM coefficients onto the cosine basis matrix.

#### Predicting stay/switch from visual cues during during 2-photon imaging

For our head-fixed 2-photon imaging setup, we modeled the relationship between visual features (female forward velocity, male-female distance, and male-female angle) by taking 2.5 seconds of cue feedback and projecting these data onto a raised cosine basis with four basis elements. We used the same model fitting and evaluation protocol described in Predicting stay/switch during natural courtship.

#### Predicting stay/switch from neural activity

We predicted stay/switch trial types during head-fixed 2-photon imaging from neural dynamics averaged from individual neuropils. For bilateral neuropils, signals were averaged across both hemispheres. To control for session-to-session variability in GCaMP expression and overall signal strength, we z-scored neural activity by subtracting the mean neural activity across all supervoxels over time and divided by the standard deviation of all supervoxels. Our pre-stimulus models were fit to neural dynamics in a four second window preceding stimulus onset (-5 seconds to -1 seconds relative to opto onset), and our post-stimulus models were fit using 4 seconds of dynamics following stimulus onset (0 to +4 seconds relative to opto onset). We used L2 regularization for all models with an inverse regularization strength set to *C* = 10*^−^*^1^. The rest of the model-fitting procedure follows Predicting stay/switch during natural courtship.

#### Tap detection model

We used a tap detector model previously described in Roemschied et al, 2023 [26]. In brief, the model was based on a convolutional neural network architecture consisting of two 2D convolutional layers followed by two fully connected layers. The outputs of each convolutional layer were passed through a RELU activation and a 2D max pooling layer with a kernel size of two and stride of two. The model was trained using the AdamW algorithm for 100 epochs with a batch size of 16 and a learning rate of 0.0001. The model was constructed and trained using the PyTorch library.

### 2-photon imaging

#### Simulated female during 2-photon imaging

A dynamic black square was used to simulate a female-like object. An LC4500 projector (Keynote Photonics) projected the image onto a spherical mirror, which reflected the image onto a plaster-coated styrofoam screen (diameter=165cm) that wrapped 270° around the fly, with the remaining 90° left open for microscope access. The display was calibrated as a full sphere, with the full projected image corresponding to 360° of azimuth and a spherical warp applied to correct for display curvature (custom Matlab script). Because the fly was positioned at the center of the sphere, the angular subtense of any stimulus was determined entirely by the projection geometry — the sphere radius cancels, as both the arc length on the screen and the distance to the screen scale proportionally. The visual stimulus’s movement consisted of two components: a stereotyped side-to-side trajectory and a naturalistic looming component played back from a courtship interaction in freely behaving animals. Since the imaged male fly’s left eye was blinded, the virtual female was presented exclusively in the right hemifield, spanning approximately 0° to 36° azimuth relative to the fly’s heading. The angular size of the stimulus on the fly’s retina was computed as *θ* = *α/d*, where *α* = 135 is a scaling constant and *d* is the male–female distance from the courtship tracking data, yielding a range of 5°–64°.

#### Hot cell stimulation during 2-photon imaging

In our 2-photon setup, optogenetic stimulation was provided by a 700 nm LED (Thor Labs M700L4) integrated into the objective. Stimulus intensity was calibrated to be consistent with the stimulation strength used in free behavior experiments. Since the HCs innervate the aristae, we focused on matching the stimulus intensity underneath the fly holder rather than optimizing the stimulus intensity directly after the objective. Underneath the fly holder (where the aristae are located), our stimulus intensity measured 0.544mW*/*cm^2^.

We provided open-loop HC stimulation in our head-fixed 2-photon imaging experiments. Inter-stimulus intervals were randomly drawn from a normal distribution with a mean time (*µ* = 29 seconds) computed from stay/switch sessions during free courtship, with a hard lower and upper bound of 15 and 45 seconds, respectively. We used 500 ms LED pulses for all experiments in this setup.

### Imaging parameters

For imaging experiments, flies were head-fixed using custom 3D-printed mounts. Following established protocols [42], the head was rotated forward 90 degrees and secured to the holder with a combination of wax and UV-curable resin (Norland Optical and Electronic Adhesives). The fly was positioned with the abdomen and thorax plane as parallel as possible to the holder to facilitate natural walking behavior. Under refrigerated saline, head cuticle, fat, and tracheal tissue were carefully removed from the brain using fine forceps.

Our functional imaging protocol acquired volumetric data at 128 × 256 pixel resolution with a 1.4 Hz sampling rate, enabling visualization of most brain regions including portions of the optic lobe. Voxel dimensions varied slightly across experiments, averaging 1.5 µm (X-dimension) × 1.5 µm (Y-dimension) × 5 µm (Z-dimension). At the conclusion of each experiment, high-resolution anatomical volumes were acquired at 512 × 1024 pixel resolution, yielding approximate voxel dimensions of 0.75 µm (X-dimension) × 0.75 µm (Y-dimension) × 2 µm (Z-dimension).

Image acquisition was controlled via Matlab using ScanImage 5.1 (Vidrio). Samples were illuminated at 920 nm using a Coherent 2-photon laser system. The laser light was directed through a long pass filter (SCHOTT RG715) and dichroic mirror positioned between the objective and detection pathways. Emitted light from the objective was separated into green and red channels using a dichroic beamsplitter (Semrock FF562-Di03), with each channel detected by independent GaAsP photo-multiplier tubes (Hamamatsu). Laser power was regulated via a pockels cell, and we applied an exponential power ramp through the pockels cell to compensate for depth-dependent signal attenuation, maintaining a maximum sample power of 20 mW. Recording sessions typically extended for 30 minutes.

### Neural Data Preprocessing

To maximize signal quality from the functional imaging data, we implemented a multi-stage preprocessing pipeline.

#### Motion Correction

Flies expressed pan-neuronal GCaMP6s for functional imaging and tdTomato as a structural marker. All functional and structural volumes were registered to a 50-volume average of the tdTomato channel, correcting for physiological motion and gradual morphological drift. Image registration was performed using ANTs [43].

#### Supervoxel Segmentation

Supervoxels were computed independently for each z-slice. We z-scored activity values and applied agglomerative clustering with spatial connectivity constraints, specifying 2000 clusters per slice. The left and right hemispheres were clustered separately and later merged.

#### Photobleaching Correction

To correct for photobleaching artifacts in the GCaMP channel, we computed a 100-sample moving average (approximately 70 seconds) within each supervoxel and subtracted it from the raw fluorescence trace.

Our imaging data contained noise from various sources, including physiological motion, shot noise, and background contamination from our display. We addressed background contamination by segmenting out non-brain voxels, averaging their fluorescence, and subtracting this signal from brain voxels.

To further eliminate residual shared noise between channels, we followed the linear regression-based approach of Hallinen et al., 2021 [44]. This method models the cleaned GCaMP signal as:

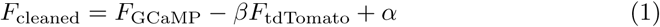

where *β* and *α* are regression coefficients that minimize the residual sum of squares:

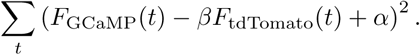

#### Signal Deconvolution

To infer neural activity from the slower calcium signal, we applied the OASIS algorithm [45] to each supervoxel’s fluorescence trace. The model assumed an autoregressive process of order 1 (AR(1)) and estimated kinetic parameters directly from the data.

### Registration to the Functional Drosophila Atlas

Our registration pipeline followed the general procedure described in Brezovec et al., 2024 [34]. First, we used ANTs to compute an affine transformation between our functional and high-resolution structural volumes. Next, a SyN warp aligned each structural volume to a common mean-brain space. Finally, we applied the BIFROST transformation pipeline to warp mean-aligned volumes into FDA (Functional Drosophila Atlas) space. All transformations were performed separately for each imaging session.

### Neural Encoding Models

To identify encoding strength for the optogenetic stimulus, we fit linear regression models (using statsmodels in Python) to each supervoxel. Predictor variables included male rotational speed, male forward speed, and the optogenetic stimulus. The optogenetic stimulus was featurized as a boxcar stimulus with amplitude 1 and spanning 5 samples (*≈* 4 seconds). We applied Benjamini–Hochberg correction to control the false discovery rate (FDR) to 5% for each session.

### Exploded brain visualizations

For maps showing raw neural activity, we registered functional volumes into the FDA anatomical space, averaged these volumes together, and took a sign-preserving maximum-projection along the anterior-posterior axis. Maps showing GLM coefficients relied on a similar procedure, but we averaged encoding model coefficients assigned to each supervoxel rather than averaging raw neural activity.

## Supplementary Information

**Fig. SI1.**
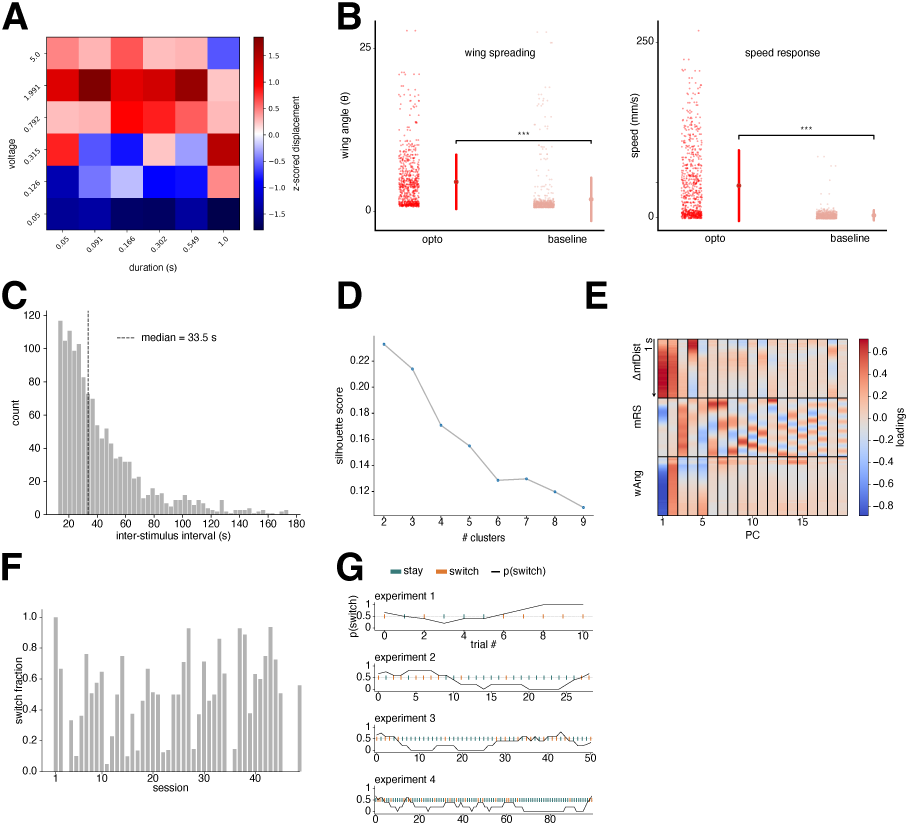
Courting male flies make stay/switch decisions in response to fictive heat threat. (A) Z-scored optogenetic stimulus-induced displacement in solitary male flies within a 0.5 post-stimulus window. Stimuli were randomly swept along a duration dimension (x axis) and voltage dimension (y axis). (B) Hot cell activation drives escape behaviors in solitary males. Left: wing spreading behavior after optogenetic stimulus (red) and at baseline (light red). Right: speed after optogenetic stimulus (red) and at baseline (light red). (C) Distribution of inter-stimulus intervals (ISIs) pooled across all courtship experiments. The median ISI is indicated by a vertical dashed line. (D) A two-cluster partitioning provides the highest silhouette score for our data (higher scores indicate better cluster assignments). (E) We reduced the dimensionality of stimulus-induced behaviors (male-female distance, male rotational speed, angle of song-producing wing) using principal components analysis (PCA). PC loadings onto original variables (y axis) are shown for the first 19 PCs (x axis), which account for *≈* 90% of the variance in these data. (F) Fraction of switch trials for all 46 experiments in our courtship dataset. (G) Empirical switch probabilities as a function of trial count for three representative courtship experiments. “Stay” trials are indicated by teal ticks, and “switch” trials are indicated by orange ticks. Black line indicates running switch probability over a five trial window.

**Fig. SI2.**
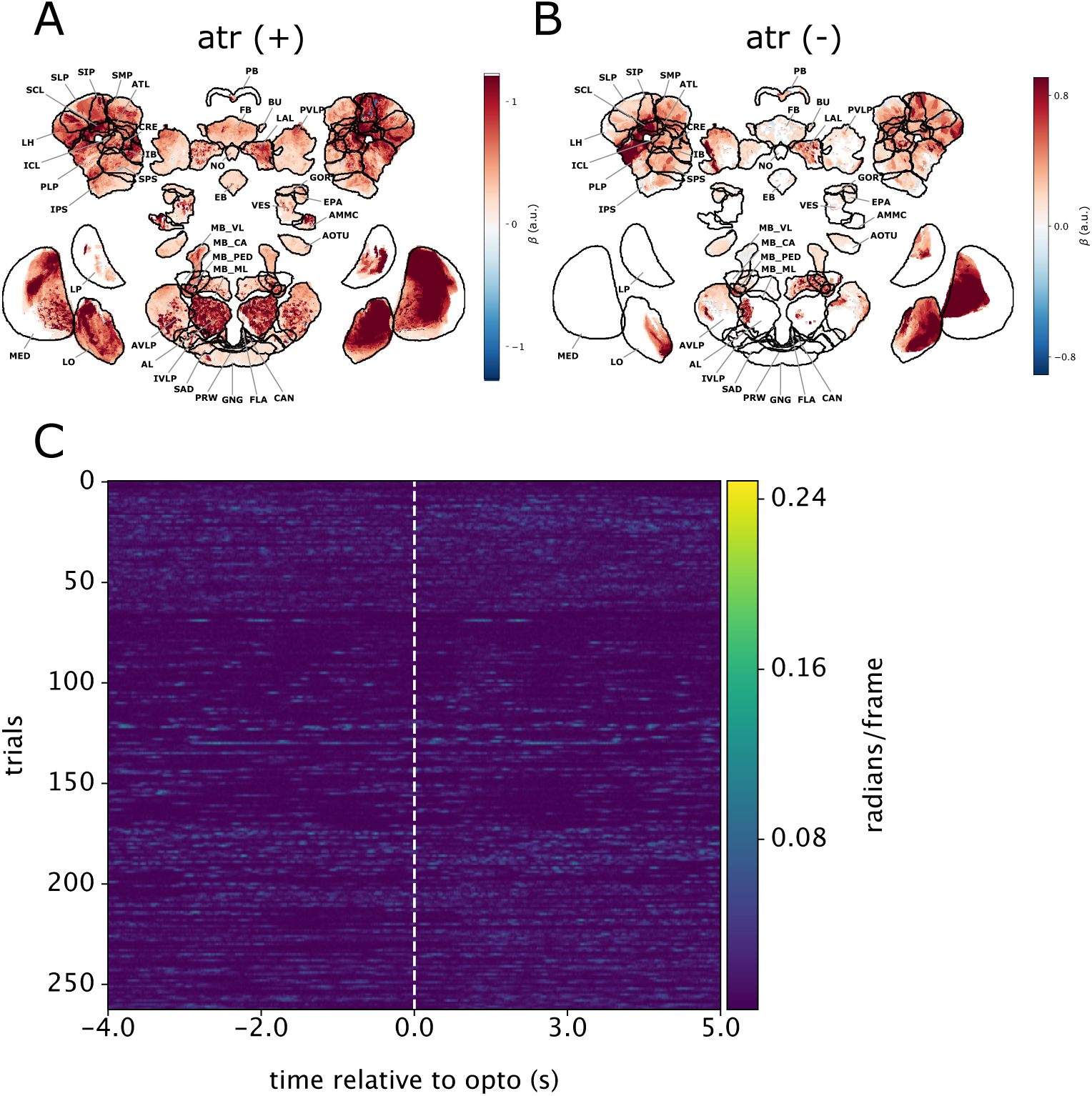
ATR-controls for stay/switch experiments in head-fixed 2-photon imaging. (A) A general linear model reconstructed neural activity from the male’s forward and rotational speed (mFS, mRS) and the optogenetic stimulus (opto). Encoding weights are shown for the optogenetic stimulus in sessions where animals were fed ATR food (n=10). (B) Same as (A), but for a fly without ATR food. (C) Male rotational speed aligned to optogenetic stimulus onset for ATR-experiments (n=4).

## Supplementary Movies

***Movie S1***

**Hot cell activation in solitary male flies** Optogenetic activation of hot cell neurons in solitary male flies using the minimal stimulus that reliably evokes escape behaviors. In this video, trials are stitched together, with each trial consisting of 1 second pre-stimulation and 1 second post-stimulation. Stimulation events are indicated with a red dot in the upper left-hand corner. This movie relates to Fig. SI1A.

***Movie S2***

**Hot cell activation in courting male flies** Optogenetic activation of hot cell neurons in male flies during courtship. Stimulation parameters are identical to those used to drive escape behaviors in solitary males. In this video, trials are stitched together, with each trial consisting of 1 second pre-stimulation and 1 second post-stimulation. Stimulation events are indicated with a red dot in the upper left-hand corner. This movie relates to Fig. 1B-C.

## References

[1] Jiang, X. & Pan, Y. Neural control of action selection among innate behaviors 38, 1541–1558. URL https://www.ncbi.nlm.nih.gov/pmc/articles/PMC9723052/.

[2] Sareen, P., Wolf, R. & Heisenberg, M. Attracting the attention of a fly 108, 7230–7235. URL https://www.pnas.org/doi/full/10.1073/pnas.1102522108.

[3] Wang, L. & Krauzlis, R. J. Visual selective attention in mice 28, 676–685.e4. URL https://www.sciencedirect.com/science/article/pii/S096098221830040X.

[4] Fernandes, A. M. et al. Neural circuitry for stimulus selection in the zebrafish visual system 109, 805–822.e6. URL https://linkinghub.elsevier.com/retrieve/pii/S0896627320309612.

[5] Bahl, A. & Engert, F. Neural circuits for evidence accumulation and decision making in larval zebrafish 23, 94–102.

[6] Skals, N., Anderson, P., Kanneworff, M., Löfstedt, C. & Surlykke, A. Her odours make him deaf: crossmodal modulation of olfaction and hearing in a male moth 208, 595–601. URL https://journals.biologists.com/jeb/article/208/4/595/9405/Her-odours-make-him-deaf-crossmodal-modulation-of.

[7] Richman, E. B., Ticea, N., Allen, W. E., Deisseroth, K. & Luo, L. Neural landscape diffusion resolves conflicts between needs across time | nature URL https://www.nature.com/articles/s41586-023-06715-z.

[8] Crickmore, M. & Vosshall, L. Opposing dopaminergic and GABAergic neurons control the duration and persistence of copulation in drosophila 155, 881–893. URL https://linkinghub.elsevier.com/retrieve/pii/S0092867413012312.

[9] Gautham, A. K., Miner, L. E., Franco, M. N., Thornquist, S. C. & Crickmore, M. A. Dopamine biases decisions by limiting temporal integration 632, 850–857.

[10] Cazalé-Debat, L. et al. Mating proximity blinds threat perception 1–9. URL https://www.nature.com/articles/s41586-024-07890-3.

[11] Coen, P. et al. Dynamic sensory cues shape song structure in drosophila 507, 233–237.

[12] Calhoun, A. J., Pillow, J. W. & Murthy, M. Unsupervised identification of the internal states that shape natural behavior 22, 2040–2049. URL https://www.nature.com/articles/s41593-019-0533-X. Number: 12.

[13] Garrett, N. & Daw, N. D. Biased belief updating and suboptimal choice in foraging decisions 11, 3417. URL https://www.nature.com/articles/ s41467-020-16964-5.

[14] Constantino, S. M. & Daw, N. D. Learning the opportunity cost of time in a patch-foraging task 15, 837–853.

[15] Stephens, D. W. & Krebs, J. R. Foraging theory Monographs in behavior and ecology (Princeton University Press).

[16] Cisek, P. Cortical mechanisms of action selection: the affordance competition hypothesis 362, 1585–1599. URL https://www.ncbi.nlm.nih.gov/pmc/articles/ PMC2440773/.

[17] Cisek, P. Evolution of behavioural control from chordates to primates 377, 20200522. URL https://royalsocietypublishing.org/doi/full/10.1098/rstb.2020.0522.

[18] Cisek, P. & Kalaska, J. F. Neural mechanisms for interacting with a world full of action choices 33, 269–298.

[19] Findling, C. et al. Brain-wide representations of prior information in mouse decision-making 645, 192–200. URL https://www.nature.com/articles/s41586-025-09226-1.

[20] Pereira, T. D. et al. SLEAP: A deep learning system for multi-animal pose tracking 19, 486–495. URL https://www.nature.com/articles/s41592-022-01426-1.

[21] Hindmarsh Sten, T., Li, R., Otopalik, A. & Ruta, V. Sexual arousal gates visual processing during drosophila courtship 595, 549–553. URL http://www.nature.com/articles/s41586-021-03714-w.

[22] Gallio, M., Ofstad, T. A., Macpherson, L. J., Wang, J. W. & Zuker, C. S. The coding of temperature in the drosophila brain 144, 614–624. URL https://www.cell.com/cell/abstract/S0092-8674(11)00067-5.

[23] Simões, J. M. et al. Robustness and plasticity in drosophila heat avoidance 12, 2044. URL https://www.nature.com/articles/s41467-021-22322-w. Number: 1.

[24] Ache, J. M. et al. Neural basis for looming size and velocity encoding in the drosophila giant fiber escape pathway 29, 1073–1081.e4.

[25] Ache, J. M., Namiki, S., Lee, A., Branson, K. & Card, G. M. State-dependent decoupling of sensory and motor circuits underlies behavioral flexibility in drosophila 22, 1132–1139.

[26] Roemschied, F. A. et al. Flexible circuit mechanisms for context-dependent song sequencing 622, 794–801. URL https://www.nature.com/articles/s41586-023-06632-1.

[27] Klapoetke, N. C. et al. Independent optical excitation of distinct neural populations 11, 338–346.

[28] McKellar, C. E. et al. Threshold-based ordering of sequential actions during Drosophila courtship 29, 426–434.e6. URL https://www.sciencedirect.com/science/article/pii/S0960982218316518.

[29] Clowney, E. J., Iguchi, S., Bussell, J. J., Scheer, E. & Ruta, V. Multimodal chemosensory circuits controlling male courtship in drosophila 87, 1036–1049.

[30] Chen, T.-W. et al. Ultra-sensitive fluorescent proteins for imaging neuronal activity 499, 295–300. URL https://www.ncbi.nlm.nih.gov/pmc/articles/PMC3777791/.

[31] Pfeiffer, B. D. et al. Tools for neuroanatomy and neurogenetics in drosophila 105, 9715–9720. URL https://pmc.ncbi.nlm.nih.gov/articles/PMC2447866/.

[32] Henry, G. L., Davis, F. P., Picard, S. & Eddy, S. R. Cell type–specific genomics of drosophila neurons 40, 9691–9704. URL https://pmc.ncbi.nlm.nih.gov/articles/PMC3479168/.

[33] Brezovec, B. E. et al. Mapping the neural dynamics of locomotion across the drosophila brain 34, 710–726.e4. URL https://linkinghub.elsevier.com/retrieve/pii/S0960982223017633.

[34] Brezovec, L. E. et al. BIFROST: a method for registering diverse imaging datasets. URL https://www.biorxiv.org/content/10.1101/2023.06.09.544408v1. Pages: 2023.06.09.544408 Section: New Results.

[35] Collie, M. F. et al. Specialized parallel pathways for adaptive control of visual object pursuit. URL https://www.biorxiv.org/content/10.1101/2025.04.23.650240v1. Pages: 2025.04.23.650240 Section: New Results.

[36] Clemens, J. et al. Connecting neural codes with behavior in the auditory system of drosophila 87, 1332–1343. URL https://www.cell.com/neuron/abstract/S0896-6273(15)00708-4.

[37] Pang, R., Baker, C. A., Murthy, M. & Pillow, J. Inferring neural population codes for drosophila acoustic communication 122, e2417733122. URL https://www.pnas.org/doi/abs/10.1073/pnas.2417733122.

[38] Tinbergen, N. The study of instinct 1st ed. 2nd impression edn (Clarendon P).

[39] Ribeiro, I. M. et al. Visual projection neurons mediating directed courtship in drosophila 174, 607–621.e18. URL https://linkinghub.elsevier.com/retrieve/pii/S0092867418307888.

[40] Brezovec, L. E., Berger, A. B., Druckmann, S. & Clandinin, T. R. Neural correlates of future volitional action in drosophila. URL https://www.biorxiv.org/content/10.1101/2023.09.08.556917v1. Pages: 2023.09.08.556917 Section: New Results.

[41] Weber, A. I. & Pillow, J. W. Capturing the dynamical repertoire of single neurons with generalized linear models 29, 3260–3289. URL https://direct.mit.edu/neco/article/29/12/3260-3289/8316.

[42] Drosophila neurobiology: a laboratory manual.

[43] Avants, B. B., Tustison, N. & Johnson, H. Advanced normalization tools (ANTS).

[44] Hallinen, K. M. et al. Decoding locomotion from population neural activity in moving c. elegans 10, e66135. URL 10.7554/eLife.66135.

[45] Friedrich, J., Zhou, P. & Paninski, L. Fast online deconvolution of calcium imaging data 13, e1005423. URL https://journals.plos.org/ploscompbiol/article?id=10.1371/journal.pcbi.1005423.

